# RNase III influences microaerobic symbiotic pathways and RNA regulation in *Sinorhizobium meliloti*

**DOI:** 10.64898/2026.02.10.705003

**Authors:** Sabina K. Guedes-García, Natalia I. García-Tomsig, Rute G. Matos, Margarida Saramago, Cecilia M. Arraiano, José I. Jiménez Zurdo

## Abstract

Bacterial ribonucleases (RNases) are central components of post-transcriptional networks underlying environmental adaptation. However, their contribution to the ecological specialization of bacteria with complex lifestyles, such as nitrogen-fixing legume symbionts, remains poorly understood. Here, we investigated the role of the double-stranded RNase III ortholog (*Sm*RNase III) in *Sinorhizobium meliloti*, the symbiotic partner of alfalfa (*Medicago sativa* L.). Loss of *Sm*RNase III function affected the expression of nearly 30% of protein-coding genes and 12% of annotated non-coding RNAs (sRNAs). Remarkably, more than 70% of these changes occurred under the microaerobic conditions typical of symbiotic root nodules. Many *Sm*RNase III-dependent transcripts encode pathways supporting microaerobic metabolism and nitrogen fixation in endosymbiotic bacteroids. Analysis of sequencing read coverage revealed putative consensus cleavage signatures biased toward mRNA 5′ untranslated regions, suggesting preferential processing at these sites. Altered expression of sRNAs and/or their predicted mRNA targets further supports a role for *Sm*RNase III in sRNA-mediated silencing. Consistently, *in vivo* and *in vitro* assays demonstrated that *Sm*RNase III is required for the repression of *nifK* (encoding the β-subunit of the nitrogenase MoFe protein) and *dctA* (encoding a major dicarboxylate transporter) by the antisense sRNA asNifK1 and the *trans*-sRNA AbcR1, respectively. Our findings reveal a major impact of *Sm*RNase III on shaping the symbiotic transcriptome of *S. meliloti* and provide a foundation for deeper investigation into the mechanisms and regulatory roles of RNase III activity in rhizobia.

## 1. Introduction

Ribonucleases (RNases) catalyze the processing and degradation of transcripts to adjust their levels to cellular needs. Consequently, these enzymes are regarded as global post-transcriptional regulators of gene expression (Arraiano et al., 2010). Bacteria are equipped with large sets of RNases with a diversity of activity mechanisms and substrate preference. Specifically, the RNase III family gathers structurally and functionally conserved endonucleolytic enzymes that cleave double-stranded RNA (dsRNA) in a strict metal-dependent manner. Eukaryotic RNase III orthologues, such as Rnt1, Drosha, and Dicer, play a central role in processing dsRNA structures to produce mature short interfering RNAs, including microRNAs (miRNAs) and small interfering RNAs (siRNAs) (Hammond, 2005). In contrast, in many bacteria RNA processing and decay are largely attributed to RNase E, an endoribonuclease that targets single-stranded RNA (ssRNA), including small regulatory RNAs (sRNAs). Meanwhile, the major role assigned to RNase III in bacteria is the maturation of ribosomal RNA (rRNA) and transfer RNA (tRNA). However, the comprehensive characterization of the prototypical member of the family, the *Escherichia coli* RNase III (*Ec*RNase III), encoded by the conserved *rnc* gene, has uncovered a broader impact of this enzyme in RNA metabolism. For example, RNase III regulates its own synthesis by removing 5’ RNA structures that preclude degradation of the *rnc* mRNA, and in a similar way it also regulates the degradation of *pnp* mRNA encoding the polynucleotide phosphorylase (PNPase) exoribonuclease (Bardwell et al., 1989; Matsunaga et al., 1996,; Régnier and Grunberg-Manago, 1990; Meur and Portier, 1992; Jarrige, 2001). It stands out that the RNase III specific activity on dsRNA substrates (*in vitro* and *in vivo*), envisages a prevalent role of RNase III as effector of mRNA processing and silencing upon base pairing with sRNAs of either the antisense (asRNAs) or *trans*-acting classes. Indeed, asRNA-mediated regulation of plasmid copy number and toxin-antitoxin systems is known to involve RNase III in some bacteria (Vogel et al., 2004; Lasa et al., 2011; Viegas et al., 2010; Svensson and Sharma, 2021; Mediati et al., 2022; Lejars et al., 2022). However, the function of RNase III in global adaptive regulatory networks has been only investigated in the model *E. coli* and a few other pathogens such as *Bordetella pertussis*, *Bacillus subtilis* or *Staphylococcus aureus* (Durand et al., 2012; Ifill et al., 2021; Lejars and Hajnsdorf, 2024). RNase III is not essential but *rnc* knockout severely hampers growth and influences virulence traits such as motility, biofilm development or host cell invasion in these bacteria.

The root nodule nitrogen (N_2_)-fixing symbiosis between soil-dwelling rhizobia and legume plants is another noteworthy example of a host-microbe interaction but with a beneficial outcome. Successful legume infection demands rhizobia to rapidly adapt to a variety of environments within the plant (Oldroyd et al., 2011). In particular, the root nodules hosting mature differentiated intracellular bacteroids provide the microaerobic conditions required by nitrogenase to reduce the inert atmospheric dinitrogen to usable ammonia (Dusha and Kondorosi, 1993; Mendoza, 1995; Patriarca et al., 2002; Yurgel and Kahn, 2008; Batista and Dixon, 2019). Therefore, symbiotic gene expression reprogramming driven by plant cues in rhizobia is expected to concern post-transcriptional regulation by both sRNAs and RNases. *Sinorhizobium meliloti*, the symbiont of alfalfa (*Medicago sativa* L.) and related legumes, is a model rhizobial species with a well-characterized non-coding transcriptome (Jiménez-Zurdo et al., 2013; Schlüter et al., 2013; Jiménez-Zurdo and Robledo, 2015; Robledo et al., 2020). We previously reported on the catalytic properties and impact on symbiosis of *S. meliloti* RNase III (*Sm*RNase III) (Saramago et al., 2018). This enzyme consists of an N-terminal catalytic domain (NucD) followed by a dsRNA-binding domain (dsRBD), both highly conserved. *Sm*RNase III cleaves efficiently dsRNA substrates *in vitro* in the presence of either Mg^2+^ or Mn^2+^ as metal co-factors. However, the optimal activity conditions for *E. coli* and *S. meliloti* RNase III such as pH and metal concentration differ significantly, suggesting a mechanistic and functional adaptation to the environments specific to the lifestyle of each bacterial species (Saramago et al., 2018; Srivastava and Srivastava, 1996; Sun, 2005). Accordingly, lack of *Sm*RNase III not only delays *S. meliloti* free-living growth but also nodulation kinetics of alfalfa plants and the onset of symbiotic nitrogen fixation.

In this work, we used strand-specific RNA-Seq to assess *Sm*RNase III-dependent alterations in the abundance of both protein-coding and non-protein coding transcripts upon *S. meliloti* growth under aerobic or microaerobic conditions. Our data unveiled a widespread influence of this RNase on the steady-state levels of mRNAs encoding proteins central to bacteroid metabolism and N_2_ fixation. Furthermore, we observed an impact of RNase III on the sRNA-mediated post-transcriptional regulation of these genes, positioning this enzyme as a key player in the riboregulatory network governing symbiotic pathways.

## 2. Materials and methodology

### 2.1. Bacterial strains, plasmids and growth conditions

The bacterial strains and plasmids used in this study along with their sources and relevant characteristics are listed in Table S1. *S. meliloti* strains were grown at 30°C in tryptone-yeast complex TY medium (Beringer, 1974). To mimic the nodule microaerobic environment, bacteria were first grown to logarithmic phase (OD_600_ of 0.4) in TY broth and microaerobiosis was then imposed by flushing flasks for 10 min with the mixture 2% O_2_/98% N_2_. Bacteria were further cultured under this condition at 60 rpm for 4 h prior to RNA extraction. *E. coli* strains were cultured in Luria-Bertani (LB) medium at 37°C. If required, antibiotics were added to the media at the following concentrations (µg/ml): tetracycline (Tc) 200, (Km) 50 for *E. coli* and 180 for *S. meliloti*.

### 2.2. Oligonucleotides

Table S2 provides the sequences and experimental applications of the oligonucleotides used in this work.

### 2.3. Construction of mutant strains

Strain Sm2020Δ*rnc* was generated by allelic exchange with pK18Δ*rnc* in Sm2020 as described (Saramago et al., 2018). Plasmids for constitutive expression of *nifK* and for IPTG-inducible expression of asRNA275 (formerly SmelA031; Schlüter et al., 2010) were generated as follows. The *nifK* coding sequence was amplified by PCR from genomic DNA of strain Sm2011 with Phusion high-fidelity DNA polymerase (Thermo Fisher Scientific) using the primers NifKFBamHI and NifKRNheI. The purified PCR product was digested with *Bam*HI/*Nhe*I, and inserted into pR-*eGFP*, yielding pR*nifK::eGFP*. To generate pSKi-asRNA275, the DNA sequence of asRNA275 and the promoter sequence *sinR-P_sinI_-TSS_sinI_* were amplified by PCR with SMa_asRNA_275F/SMa_asRNA_275XbaI and sinR_NdeIF/TSS3_28pb_b_sinR primer pairs. These overlapping amplification fragments were jointly used as the template for a second PCR with primers sinR_NdeIF and SMa_asRNA_275XbaI. The final PCR product was restricted with *Nde*I and *Xba*I, and then inserted into pSRKKm. Plasmid constructs were verified by Sanger sequencing. Both plasmids, pSKi-asRNA275 and pR*nifK::eGFP*, were mobilized to Sm2020 and Sm2020Δ*rnc* strains by biparental mating. Similarly, plasmid pSKiAbcR1, which drives IPTG-induced expression of the AbcR1 sRNA (García-Tomsig et al., 2022), was conjugated into the same strains to assess AbcR1- and *Sm*RNase III-dependent endogenous *dctA* levels.

### 2.4. RNA isolation

Total RNA was extracted from bacterial cultures using acid phenol-chloroform as described (Del Val et al., 2007). RNA samples were further treated with Invitrogen TURBO^TM^ DNase for 1h at 37°C, and subsequently cleaned up with the RNeasy Mini Kit (Qiagen) following the manufacturer guidelines. To enhance retention of the small RNA fraction, samples were mixed with seven volumes of 100% EtOH prior to column loading.

### 2.5. RNA sequencing and data analysis

RNA samples derived from wild type and SmΔ*rnc* strains grown under aerobic and microaerobic conditions (three biological replicates per strain and growth condition) were used to generate strand-specific cDNA libraries that were sequenced on the Illumina NextSeq Mid 500 platform to render paired-end reads (2 x 75 pb). Data were processed using R software (https://www.r-project.org/). Demultiplexed sequencing reads were mapped to the *S. meliloti* Rm1021 reference genome with the Rbowtie2 package v2.2.3 using standard parameters (Shlüter et al., 2013; Au et al., 2010). Optical duplicates were removed with SAMtools (Li et al., 2009). Uniquely mapped reads were assigned to protein coding genes or non-coding RNAs with the Rsubread package v3.12 (Liao et al., 2019). Differential gene expression analysis was conducted with the DESeq 2 package v1.34.0 (Love et al., 2014). Coverage and transcriptome-wide read distributions were visualized with the Integrative Genomic Viewer (IGV) v2.12.3 software (Robinson et al., 2011). Plots were generated using the ggplot2 package v3.3.6 (Wickham, 2009). Searches for conserved motifs were done with the MEME algorithm (http://meme-suite.org/index.html). Venn diagrams were generated with the Venny 2.0 tool (https://bioinfogp.cnb.csic.es/tools/venny/index2.0.2.html). IntaRNA (v 3.2.0; http://rna.informatik.uni-freiburg.de/) was used to predict sRNA-mRNA base-pairing interactions. RNA secondary structures were predicted using RNAfold from the ViennaRNA package (Lorenz et al., 2011).

### 2.6. RT-qPCR

cDNA was synthesized with the Takara Prime Script RT Master Mix using 1 μg of total RNA. RT-qPCR was performed on a QuantStudio 3 system (Thermo Fisher Scientific) using the Takara TB Green Premix ExTaqII with 0.5 µl of cDNA. Transcript abundance ratios were calculated as the average ΔΔ-CT (where CT is the threshold cycle) of the results of three amplifications using three independent RNA extracts, where ΔΔ CT represents the level of gene expression in the IPTG-induced strain relative to the untreated control strain. The seemingly constitutive gene *SMc01852*, encoding a phosphofructokinase, was used to normalize gene expression (Becker et al., 2004). Non-reverse transcriptase (-RT) control reactions on the RNA samples were performed simultaneously to confirm the absence of DNA contamination.

### 2.7. In vitro activity assays

The plasmid encoding His-tagged *Sm*RNase III was transformed into *E. coli* BL21(DE3) recArnc105 strain (Amarashinghe et al., 2001). The expression and purification of the recombinant S*m*RNase III was performed as previously described (Saramago et al., 2018). DNA templates for *in vitro* transcription of *Sm*RNaseIII substrates were generated by PCR using genomic DNA from *S. meliloti* Sm2011 strain (primer pairs on Table S2). The phage T7 RNA polymerase promoter sequence was incorporated into the forward primers. *In vitro* transcription was carried out using the TransGen Biotech T7 High Efficiency Transcription Kit by overnight incubation at 37°C. The DNA template was removed by incubation of the reactions 30 min at 37°C in the presence of 1 U DNase. RNA transcripts were purified by electrophoresis on an 8.3 M urea/6% polyacrylamide gel. Gel slices containing the RNA transcripts were eluted overnight at room temperature with elution buffer [0.5 M ammonium acetate pH 5.2, 1 mM EDTA, 2.5% (v/v) phenol pH 4.3]. The RNA was ethanol-precipitated, resuspended in RNase free water and quantified using the Invitrogen Qubit^TM^ RNA HS Assay Kit. The RNA substrates as275 and *nifK* mRNA were labeled at their 5′ ends with [γ-^32^ATP] using New England Biolabs T4 Polynucleotide Kinase in a standard reaction. Labeled substrates yields (cpm/μl) were determined by scintillation counting. The hybridization between labeled and non-labeled substrates to generate dsRNA was always performed in 30 mM Tris-HCl solution (pH 8) by incubation for 10 min at 80°C, followed by 45 min at 37°C. The molar ratio of unlabeled to labeled substrate molecules was 5:1.

*Sm*RNaseIII activity assays were performed in a final volume of 50 μl containing the activity buffer (30 mM Tris-HCl pH 8, 160 mM NaCl, 10 mM MgCl_2_ and 0.1 mM DTT) and the RNA substrate (concentrations indicated in the figure legends). As a control, mock-treated reactions were incubated until the end of the assay. The reactions were initiated by the addition of the enzyme, and further incubated at 30°C. At designated time points, 5-μl aliquots were withdrawn and mixed with formamide containing dye supplemented with 10 mM EDTA. Reaction products were resolved in a 7M urea/polyacrylamide gel (polyacrylamide concentrations are indicated in the respective figure legends) and visualized by digital imaging of the exposure plate (Fuji), on a Personal Molecular Imager FX scanner and analyzed with the Quantity One software (Bio-Rad). Each activity assay was performed at least in triplicate.

## 3. Results

### 3.1. SmRNase III-dependent alterations of the S. meliloti transcriptome

Lack of *Sm*RNase III confers pleiotropic free-living and symbiotic phenotypes to *S. meliloti* (Saramago et al., 2018), suggesting a broad influence of this enzyme on the turnover of the cellular transcripts. We have therefore investigated the differences in the transcriptomic profiles between a wild-type strain (Sm2011) and its *rnc* deletion mutant derivative (SmΔ*rnc*) when cultured under the aerobic and microaerobic (2% O_2_) conditions prevailing during *S. meliloti* free-living growth and symbiotic N_2_-fixation, respectively. Illumina-based sequencing of rRNA-depleted bacterial RNA delivered an average of 10 million reads per sample, which we mapped to the *S. meliloti* Rm1021 reference genome retrieved from the RhizoGATE database (Becker et al., 2009). This annotation incorporates the *trans*-sRNA (605) and asRNA (3,720) *loci* identified experimentally by genome-wide determination of transcription start sites (Schlüter et al., 2010; Schlüter et al., 2013). After normalization of read counts, only loci with coverage exceeding 10 reads in at least two of the experimental conditions were retained for further analysis. Genes were considered differentially expressed between strains and/or growth conditions if they exhibited at least a 2-fold change in read counts (*i.e.*, log_2_FC ≥1) with a *P* value ≤0.05 across the three biological replicates (Dataset S1 S3). These replicates showed a high degree of concordance in gene expression (R > 0.95), as revealed by multidimensional scaling (MDS) and by clustering of the top 250 differentially expressed-genes (Fig S1A), thus confirming the reliability and robustness of our transcriptomic dataset. Besides, the absence of reads mapping to the *rnc* locus in the mutant strain (Fig S1B), along with the upregulation of *bona fide* N_2_-fixation genes in Sm2011 under microaerobic conditions (Fig S1C), further validated the experimental setup. As independent experimental validation, RT-qPCR confirmed upregulation of *nifA* and *fixK1* in both the wild-type strain and the SmΔ*rnc* mutant, specifically under microaerobic conditions (Fig S1D).

Our analysis evidenced that microaerobiosis imposed to Sm2011 log cultures drove differential expression of 796 out of the ∼6,200 protein-coding genes (mRNAs) predicted in the genome (∼13%), roughly confirming previous microarray data from similar experiments (Becker et al., 2004) (Fig. 1A). Additionally, we also identified 97 microaerobiosis-dependent sRNAs, 37 and 60 of those annotated as *trans*-acting and asRNAs, respectively, all of them with an unknown function. *Sm*RNase III knockout caused misregulation of 654 mRNAs (10.5% of the genome) in aerobic conditions, whereas upon microaerobic growth this number raised to 1,706 (27%). Microaerobiosis similarly boosted differential expression of sRNAs in SmΔ*rnc*, increasing their number from 96 (40 *trans*-sRNAs/56 asRNAs) identified in aerobically-growth bacteria to 300 (118 *trans*-sRNAs /182 asRNAs) (Dataset S1). It is also noteworthy that a substantial number of leader mRNAs, 302 in total, are differentially expressed in the absence of *Sm*RNase III under both oxic and microoxic culture conditions (Fig 1A). Volcano plots illustrating the scattering of differentially expressed mRNAs and sRNAs in SmΔ*rnc* when compared to the wild-type strain showed that 524 of those were downregulated by the loss of *Sm*RNase III activity, whilst 388 were upregulated during bacterial growth in aerobic conditions. Under microaerobiosis, lack of *Sm*RNase III negatively affected accumulation of 1,156 transcripts, whereas the levels of 1,186 increased (Fig 1B). Notably, in this condition the most significantly affected genes mapped to the *S. meliloti* symbiotic megaplasmid pSymA. Further comparative analysis of our transcriptomic datasets, illustrated through Venn diagrams, revealed that the absence of *Sm*RNase III alone is sufficient to cause misregulation of 1,606 mRNAs and 537 sRNAs across both tested growth conditions. This corresponds to approximately 25% of the protein-coding and 12% of the non-coding transcriptome of *S. meliloti*, indicating that *Sm*RNase III exerts a broad regulatory influence, either directly or indirectly (Fig. 1C). Notably, over 70% of these transcripts (1,132 mRNAs and 393 sRNAs) are specifically affected under microaerobic conditions. These findings underscore the pivotal role of *Sm*RNase III in maintaining *S. meliloti* RNA homeostasis, particularly under the microoxic conditions characteristic of symbiotic root nodules.

**Fig. 1.**
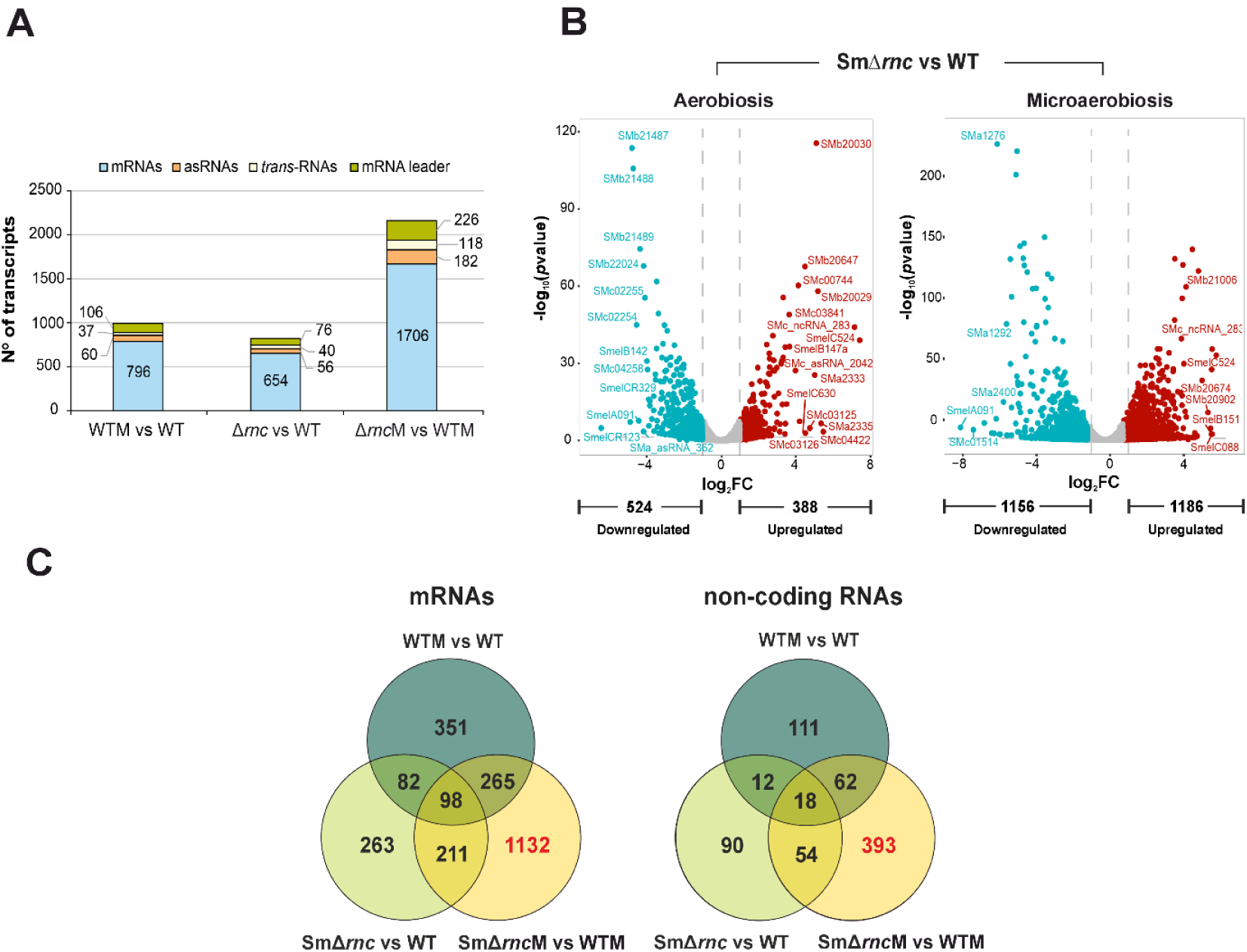
Overview of the RNA-Seq data. A) Number of differentially expressed mRNAs, asRNAs, *trans*-sRNAs, and mRNA leaders identified in each transcriptomic comparison. M, microaerobic conditions; WT, wild-type strain; Δ*rnc*, *Sm*RNase III deletion mutant (SmΔ*rnc*). B) Volcano plots showing the log₂ fold change (x-axis) versus the –log₁₀ adjusted *P*-value (y-axis) for each gene in the SmΔ*rnc* vs WT comparison in aerobic and microaerobic conditions as indicated on top. Genes with statistically significant differential expression (*P* < 0.05) are highlighted in color: upregulated genes in red and downregulated genes in blue. Genes without significant differential expression are shown in gray. Horizontal and vertical dashed lines indicate the significance threshold (*P* < 0.05) and fold change cutoff (log₂FC ≥ 1), respectively. C) Venn diagrams comparing differentially expressed mRNAs and sRNAs across transcriptomic comparisons. Transcripts uniquely misregulated in the SmΔ*rnc* mutant compared to the wild-type strain are considered *Sm*RNase III-dependent. Numbers of misregulated transcripts in the mutant and specifically under microaerobic conditions are highlighted in red.

### 3.2. Widespread effect of SmRNase III on genes involved in iron uptake, N_2_ fixation and microaerobic metabolism

To identify the cellular processes affected by the absence of *Sm*RNase III in *S. meliloti*, we performed a gene ontology (GO) analysis on the RNA-Seq datasets (Alexa and Rahnenfuhrer, 2006) (Fig. 2). *Sm*RNase III-dependent genes with predictable function, and identified under aerobic and microaerobic growth conditions were categorized into 72 and 82 established GO terms (P<0.05), respectively (Dataset S1). A significant number of genes with altered expression in the SmΔ*rnc* mutant are involved in biosynthetic and energy metabolism pathways. A notable subset of these genes is responsible for the uptake of iron complexed with the siderophore rhizobactin 1021, which is specifically secreted by the *S. meliloti* 1021/2011 strains (Persmark et al., 1993). The oxygen downshift promoted upregulation of the whole set of regulatory and structural genes involved in rhizobactin 1021 biosynthesis and transport in the wild-type strain. However, in the SmΔ*rnc* mutant this iron uptake pathway was markedly silenced under both aerobic and microaerobic growth conditions, suggesting that *Sm*RNase III positively regulates these genes at the post-transcriptional level in an O_2_-independent manner.

**Fig. 2.**
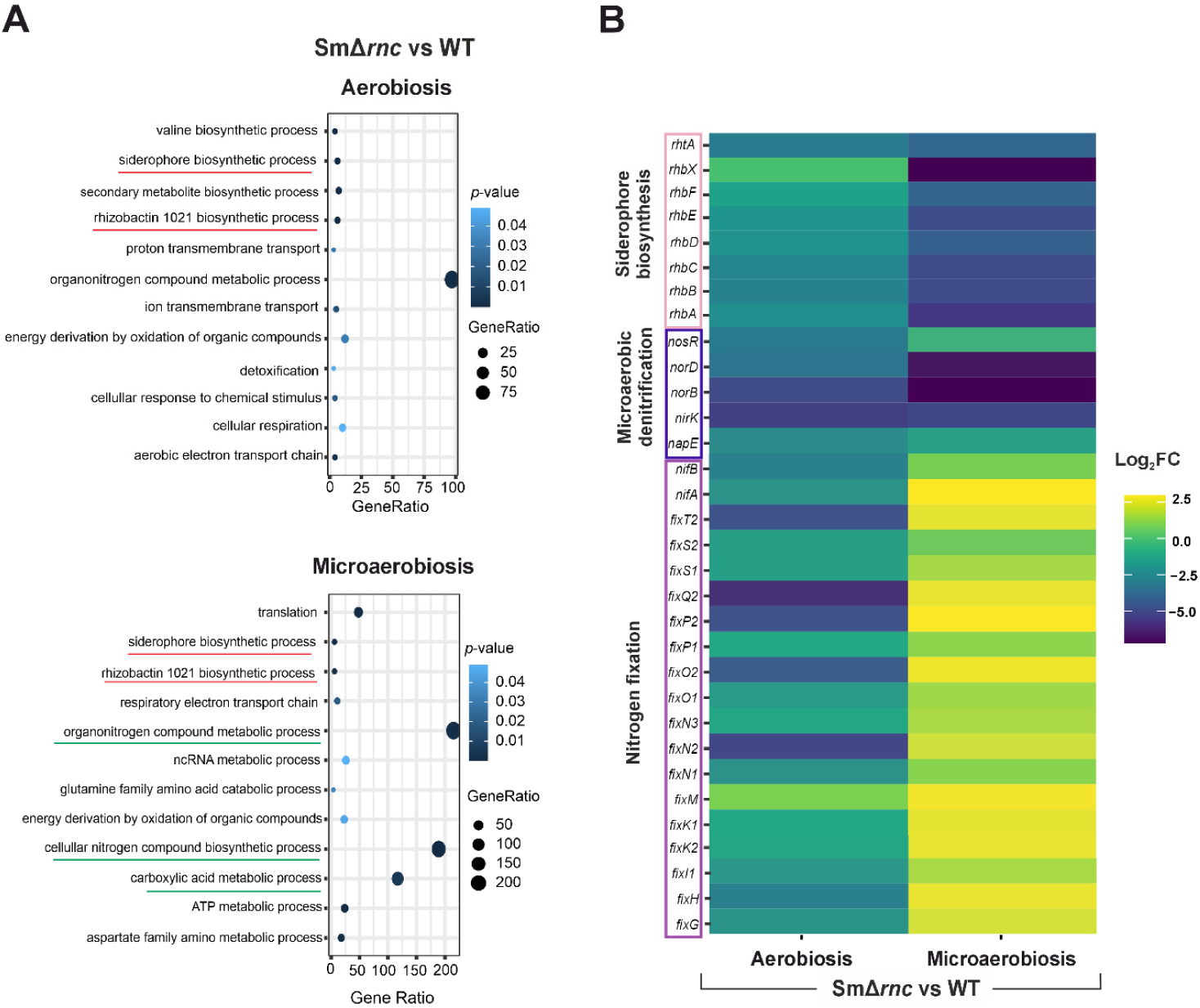
Gene ontology (GO) analysis of *Sm*RNase III-dependent mRNAs. A) Dot plot showing misregulated pathways in the SmΔ*rnc* mutant under aerobic and microaerobic conditions. Pathways highlighted in red are downregulated, while those in green are upregulated. Dot size reflects the number of genes associated with each GO term, and color intensity indicates statistical significance (*P*-value as indicated to the right). B) Heatmap of differentially expressed genes involved in iron uptake, microaerobic denitrification, and N_2_ fixation pathways. Color intensity represents relative expression levels, with yellow indicating higher expression and dark green lower expression in the SmΔ*rnc* mutant compared to the wild-type strain (WT).

*Sm*RNase III also influenced the expression of numerous genes known to support bacteroid metabolism within root nodules, most notably in microaerobic cultures. These include some involved in N_2_ fixation, microaerobic N metabolism or dicarboxylate transport. The levels of mRNAs encoding the master transcriptional regulators of symbiotic nitrogen fixation, *nifA* and *fixK1/2*, increased in the SmΔ*rnc* mutant, indicating negative post-transcriptional regulation by *Sm*RNase III. Of note, we found a similar *Sm*RNase III-dependent expression pattern for the two copies of the FixK-dependent gene cluster *fixNOQP*, required for the synthesis of the cbb3 cytochrome oxidase. Denitrification was another noteworthy microaerobic metabolic pathway influenced by *Sm*RNase III. Specifically, *napE*, *nirK*, and the gene cluster *norECBQD*, which encode periplasmic nitrate reductase, copper nitrite reductase and nitric oxide reductase, respectively, were all downregulated in Sm*Δrnc*, more significantly in microaerobiosis. Overall, these findings predict a major impact of *Sm*RNase III in the post-transcriptional fine-tuning of bacteroid metabolism.

### 3.3. Inferring putative SmRNase III substrates from the RNA-Seq data

The misregulated mRNAs and sRNAs in SmΔ*rnc* are either direct *Sm*RNase III substrates or indirectly influenced secondary targets. A detailed analysis of sequencing coverage across annotated genomic features may reveal transcript-processing signatures, thereby facilitating the identification of potential endogenous substrates of *Sm*RNase III. In addition to its own mRNA (*rnc*), bacterial RNase III is well known for processing *pnp* mRNA, as well as rRNA and tRNA precursors (Jarrige, 2001; Meur and Portier, 1992). In our experiments, *pnp* and 23S rRNA accumulated in SmΔ*rnc* as 5’-extended isoforms of the respective wild-type mature RNA species (Fig. 3). Besides, SmΔ*rnc*-derived reads corresponding to several tRNA genes (e.g., tRNA^Ser^) mapped also beyond the annotated 3’-ends of these transcripts. These coverage patterns suggest that *Sm*RNase III cleaves *S. meliloti pnp*, rRNA and tRNA precursors at either their 5’ or 3’regions, as described in other bacterial species. A genome-wide search for elongated mRNA isoforms with significantly increased accumulation in their virtual 5’-untranslated regions (UTR; extending from nucleotide positions −50 to +100 relative to the translation start codon) in SmΔ*rnc*, compared to the wild-type, identified a list of 141 and 585 *Sm*RNase III possible target candidates in aerobiosis and microaerobiosis, respectively. This list includes the *rnc* mRNA (the 5’-UTR of the *rnc* gene was not deleted in the mutant) a conserved known RNase III substrate in bacteria (Gordon et al., 2017; Matsunaga et al., 1996).

**Fig. 3.**
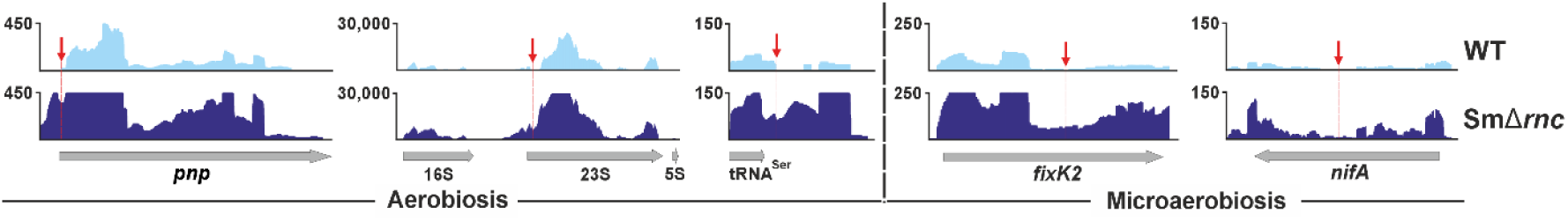
Read coverage profiles of putative endogenous *Sm*RNase III substrates in the wild-type strain (WT) and the SmΔ*rnc* mutant. Culture conditions are indicated below each panel. The height of the coverage tracks reflects the relative abundance of transcripts. The red arrow marks a specific region where a sharp drop in read density is observed, consistent with a potential *Sm*RNase III cleavage site.

RNase III-mediated cleavage may also occur within the mRNA CDS or internal to sRNAs at sites that are significantly enriched in the SmΔ*rnc* mutant with respect to the wild-type strain, where they may even be uncovered. This was for example evident in the *fixK2* and *nifA* mRNAs (Fig. 3). Scanning of the *S. meliloti* genome in 20-nt windows to search for this coverage hallmark unveiled 572 mRNAs and 72 sRNAs as additional *Sm*RNase III target candidates. To investigate potential structural or sequence-based cleavage signatures we performed an analysis in which 100-nt sequences flanking the predicted cleavage sites were extracted and compared. Based on structural similarity determined by RNAfold predictions, the sequences were grouped into four distinct structural clusters. Structural conservation among cluster members was assessed through a dot-bracket heatmap (Fig. S2). This approach enables the inference of conserved structural control points that may be functionally relevant and potentially involved in RNA processing by *Sm*RNase III as described (Gordon et al., 2017; Huang et al., 2020). Although RNase III primarily recognizes structural features, it also exhibits moderate preferences for specific local sequences at cleavage sites, such as AU-rich regions flanking the site and GC-rich sequences in the resulting 2-nucleotide overhangs, which may influence the efficiency and precision of RNA processing. (Zhang and Nicholson, 1997; Huang *et al*., 2020). To identify such potential consensus sequences, we analyzed each cluster individually. No conserved motifs were found in structural Clusters I, II, and III. However, Cluster IV consistently displayed the conserved motif ‘UCGGCAU’ across all included sequences, with an E-value of 1.4 × 10⁻³ (Fig. S2).

To evaluate whether *Sm*RNase III preferentially cleaves 5′-UTR regions, we compared predicted cleavage sites with annotated virtual 5′ UTRs (Dataset S2). A chi-squared test revealed a significant preference for cleavage within 5′-UTRs, suggesting that *Sm*RNase III modulates transcript stability or maturation at this level. Predicted *Sm*RNase III substrates are enriched in mRNAs involved amino acid catabolism, nucleotide metabolism, and gene regulation. Notably, pathways related to the degradation of glutamine family and arginine-derived amino acids, as well as cyclic nucleotide biosynthesis, were also significantly enriched, highlighting a potential role in N balance and metabolic signaling. Additionally, enrichment of transcriptional and RNA biosynthetic processes suggests the *Sm*RNase III may regulate gene expression at both transcriptional and post-transcriptional levels. These findings support a model in which *Sm*RNase-mediated RNA processing is tightly integrated with cellular metabolic reprogramming and stress adaptation.

### 3.4. SmRNase III regulation of mRNA turnover induced by antisense transcription

Transcriptomic analysis of SmΔ*rnc* unveiled a total of 56 and 182 asRNAs differentially expressed after aerobic and microaerobic bacterial growth, respectively, with 20 common to both conditions (Fig. 1A). Notably, among these transcripts are SMc_asRNA_744, SMc_asRNA_1551, SM_asRNA_1120, SMa_asRNA_380, and SMc_asRNA_936, which are antisense to the protein-coding genes *metB*, *dgkA*, *dgkB*, *napC*, and *cyaE1*, respectively. These genes are involved in fundamental metabolic and regulatory pathways, including methionine biosynthesis, membrane lipid turnover, periplasmic nitrate reduction, and nucleotide signaling.

Additionally, 36 and 162 asRNAs were misregulated in the absence of *Sm*RNase III under aerobic and microaerobic conditions, respectively. Notably, nearly 90% (142) of the microaerobically misregulated asRNAs were upregulated in the mutant strain, including those antisense to key N_2_ fixation genes such as *nifK* (encoding a nitrogenase structural component), *nifN* (involved in Fe-Mo cofactor biosynthesis), and *fixX* (encoding a ferredoxin). The asRNA SMc_asRNA_1268, antisense to *nouN*, was also induced in the *Sm*RNase III mutant in microaerobiosis. NouN is a component of the NADH-ubiquinone oxidoreductase complex, a central enzyme in aerobic respiration and energy production. Collectively, these findings suggest a prominent role for the concerted activity of asRNAs and *Sm*RNase III in fine-tuning aerobic energy metabolism and microaerobic N_2_ fixation in endosymbiotic bacteroids.

### 3.5. SmRNase III mediates nifK silencing driven by asnifK1

To further investigate the role of *Sm*RNase III as an effector of mRNA silencing mediated by asRNAs, we performed a detailed analysis of the interaction between *nifK* mRNA and asRNA275. The *nifK* gene (*SMa0829*) encodes the β-subunit (FeMo protein) of the *S. meliloti* nitrogenase complex. Among the three sRNAs likely transcribed antisense to *nifK*, asRNA275, also annotated as SMa_asRNA_275 or SmelA031, was prominently represented in our RNA-Seq dataset, showing strong upregulation in the SmΔ*rnc* mutant under microaerobic conditions (Fig. 4A). Accordingly, we designated this transcript as asnifK1. To assess *nifK* regulation by asnifK1, we employed a dual-plasmid assay in which the asRNA and its mRNA target were expressed independently from compatible plasmids, bypassing their native promoters. To confirm IPTG-induced overexpression of asnifK1 from plasmid pSKi-asnifK1, we first performed RT-qPCR in the wild-type strain grown aerobically in TY medium (Fig. 4B, left). Under these conditions, abundance of *nifK* mRNA, constitutively expressed from a synthetic promoter (*P*syn) on plasmid pR-*nifK*, was significantly reduced in wild-type cells overexpressing asnifK1 following IPTG induction, but remained unchanged in the SmΔ*rnc* mutant (Fig. 4B, right). These results provide the first genetic evidence supporting the involvement of *Sm*RNase III in asnifK1-dependent silencing of *nifK* mRNA.

**Fig. 4.**
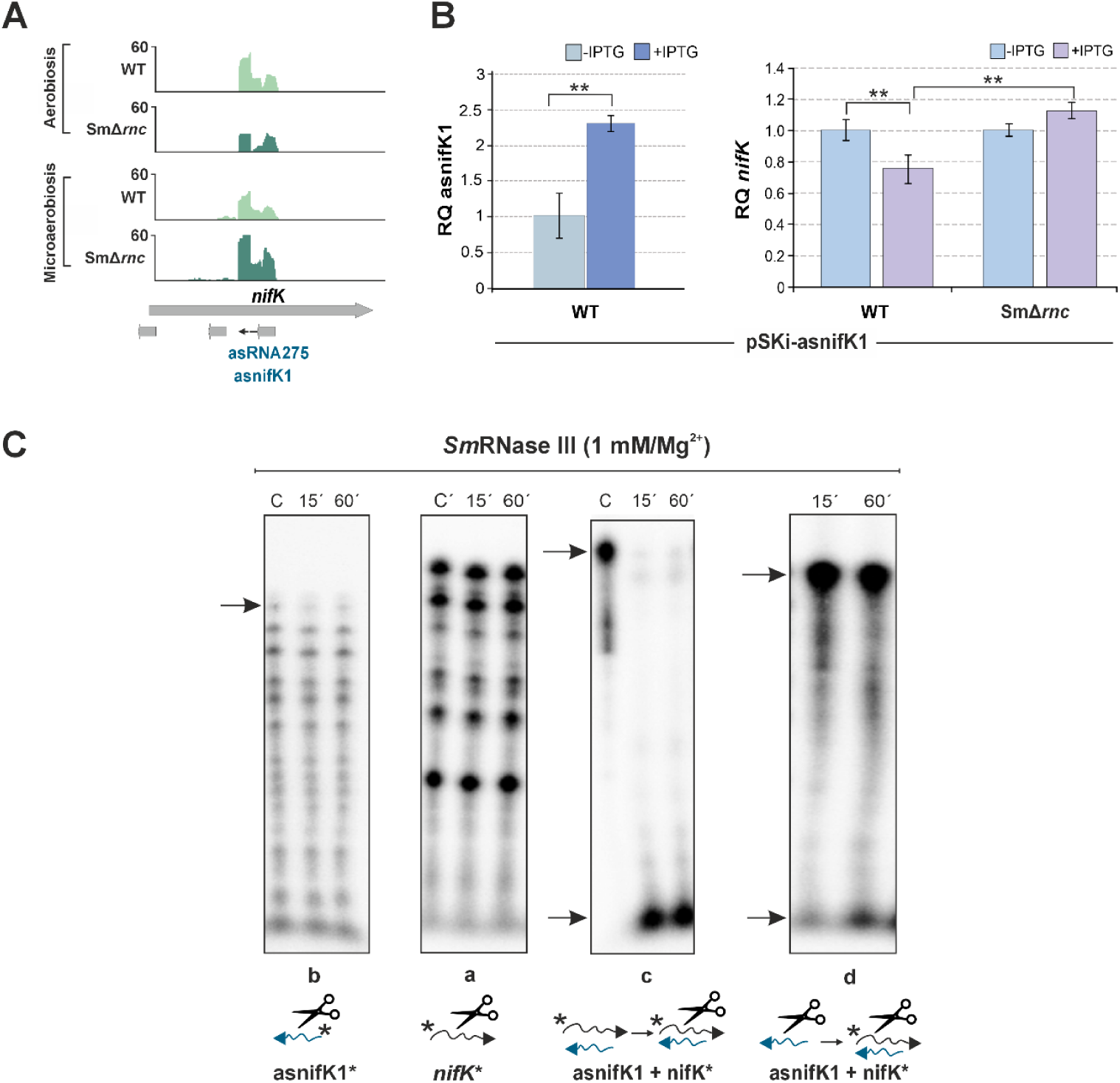
*Sm*RNase III-assisted asnifK1-mediated regulation of *nifK*. A) Read coverage of asnifK1 in wild-type (WT) and SmΔ*rnc* strains under aerobic and microaerobic conditions. B) RT-qPCR analysis of the impact of ectopic IPTG-induced asnifK1 overexpression from plasmid pSKi-asnifK1 on *nifK* mRNA abundance. Determinations were performed in total RNA extracted from Sm2020 cells transformed with pSKi-asnifK1 and cultured in TY medium to exponential phase. Three independent biological replicates were obtained, and each was split into IPTG-treated and untreated subcultures and incubated for an additional 4 hours. For each biological replicate, RT-qPCR determinations were carried out in duplicate (n=6). Relative quantification (RQ) values were normalized to *SMc01852* as a constitutive control. Asterisks indicate statistically significant differences according to ANOVA (P < 0.05); NS indicates non-significant differences. C) *Sm*RNase III activity on asnifK1, *nifK*, and the duplex formed between them. *In vitro* reactions were performed at 30 °C for the time points (minutes) indicated above each panel. “C” denotes control reactions without enzyme. The wild-type enzyme (1 mM) was incubated with 1 pmol/µl of RNA substrates in the presence of 10 mM MgCl₂. Reaction products were analyzed on 7 M urea/6% polyacrylamide gels. Arrows indicate RNA depletion in reactions containing the enzyme compared to controls. The order in which the enzyme and the RNA substrates were added is shown below each panel and asterisks indicate the radiolabeled RNA.

We next performed *in vitro* cleavage assays with *nifK* and asnifK1 transcripts using purified *Sm*RNase III (1 µM) and Mg²⁺ as a cofactor (Fig. 4C). The enzyme was first incubated with radiolabeled *nifK* and asnifK1 transcripts in separate reactions. These assays revealed modest cleavage of asnifK1 by *Sm*RNase III, as evidenced by time-dependent degradation of the full-length transcript (highlighted with an arrow), whereas *nifK* remained unaffected (Fig. 4C, panels a and b). However, when radiolabeled *nifK* was incubated with unlabeled asnifK1 to promote duplex formation, the substrate was completely degraded after 15 minutes of incubation with *Sm*RNase III (Fig. 4C, panel c). To test whether prior processing of the asRNA affects duplex cleavage, asnifK1 was pre-incubated with *Sm*RNase III before the addition of radiolabeled *nifK*. The results showed that *nifK* degradation still occurred, albeit with significantly reduced efficiency (Fig. 4C, panel d).

Together, these findings indicate that asnifK1 is an endogenous substrate of *Sm*RNase III and that antisense pairing enhances *nifK* decay by the enzyme.

### 3.6. SmRNase III assists regulation by trans-sRNAs

Bacterial RNase III also functions as a silencing enzyme of mRNAs that base-pair with *trans*-encoded sRNA regulators. In *S. meliloti*, the only mRNA targetomes characterized genome-wide are those of the homologous *trans*-sRNAs AbcR1 and AbcR2, which act as central regulatory hubs within extensive metabolic networks (García-Tomsig et al., 2022). The steady-state levels of AbcR1 remained unaffected by microaerobic conditions or deletion of the *rnc* gene, indicating that *Sm*RNase III does not participate in its turnover (Dataset S1). Nonetheless, 111 and 146 *Sm*RNase III-dependent mRNAs are part of the AbcR1 and AbcR2 regulons, respectively (Dataset S3; Fig. 5A). Functional analysis of these mRNA sets revealed enrichment in pathways related to carbohydrate metabolism, environmental stress response, microaerobic adaptation, N_2_ fixation, redox homeostasis, and the transport of various nutrients, including C4-dicarboxylates (Dataset S3).

**Fig. 5.**
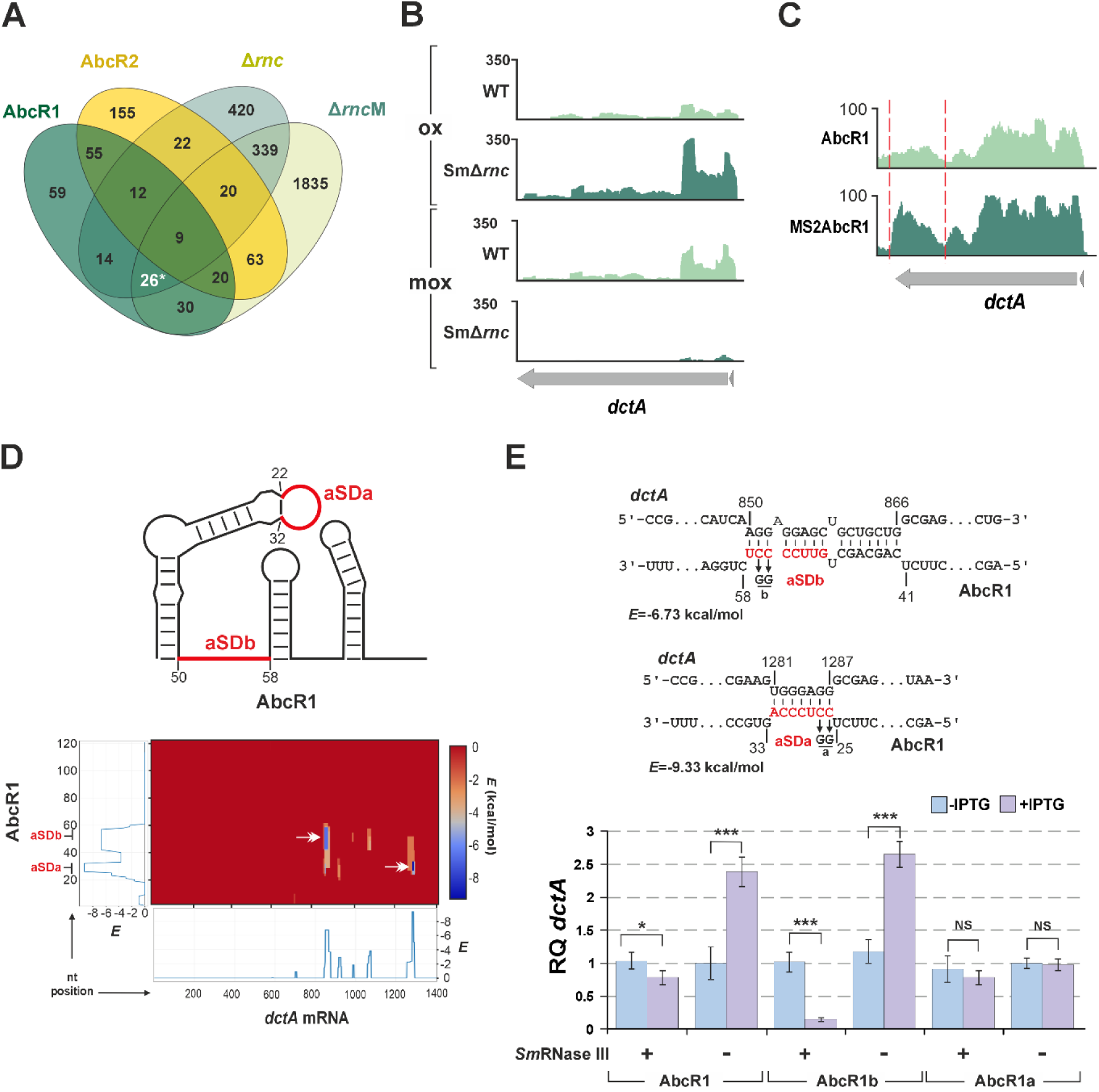
*Sm*RNase III-assisted *trans* regulation of *dctA* by AbcR1 A) Venn diagram comparing the AbcR1 and AbcR2 mRNA interactomes with the sets of mRNAs differentially accumulated in the SmΔ*rnc* mutant under aerobic and microaerobic (M) conditions. The subset marked with an asterisk includes *dctA*. B) Read coverage profile of *dctA* in wild-type (WT) and SmΔ*rnc* strains grown under aerobic (ox) and microaerobic (mox) conditions. C) Recovery profiles of *dctA* mRNA from affinity chromatography using MS2-tagged AbcR1 as bait. Wild-type AbcR1 served as a specificity control. The enriched mRNA region is indicated by dashed lines. D) IntaRNA prediction of the base-pairing profile between AbcR1 and *dctA*. The schematic at the top depicts the predicted AbcR1 secondary structure, highlighting the aSDa and aSDb regulatory motifs in red. The plot below shows position-wise minimal energy profiles for *dctA* and AbcR1, along with a heatmap of predicted interactions. Axes indicate nucleotide positions along each RNA. Interaction patterns highlight potential binding regions that may mediate regulatory pairing. Arrows mark the wild-type interactions between aSDa/aSDb and *dctA*. E) Predicted base-pairing interactions between the aSDa and aSDb motifs of AbcR1 and *dctA* mRNA are shown (upper panel). The minimum hybridization energy (*E*) is indicated. Nucleotide positions are given relative to 50 nt upstream of the *dctA* start codon and the transcription start site of AbcR1. Nucleotides in red correspond to targeting motifs, and point mutations in each regulatory motif are indicated. Lower panel, RT-qPCR analysis of the impact of IPTG-induced overexpression of AbcR1, AbcR1a, and AbcR1b (from pSKi derivatives) on *dctA* mRNA abundance in both the wild-type strain (+) and the SmΔ*rnc* mutant (-). Total RNA was extracted from Sm2020 cells transformed with the corresponding pSKi derivative after growth in TY medium to exponential phase. Three independent biological replicates were split into IPTG-treated and untreated subcultures and incubated for an additional 4 hours. Relative quantification (RQ) values were normalized to *SMc01852* as a constitutive control. Plotted values correspond to two determinations in RNA derived from each biological replicate (n=6). Asterisks indicate statistically significant differences according to ANOVA (*p < 0.05; **p < 0.01; ***p < 0.001); NS indicates non-significant differences.

Bacteroids use dicarboxylic acids as preferred carbon sources to meet the high energy demands of N_2_ fixation. *Sm*RNase III influences *dctA*, a key dicarboxylate transporter mRNA, in a condition-dependent manner (Fig. 5B). Under aerobic conditions, *dctA* is upregulated in the SmΔ*rnc* mutant, whereas its levels decrease in microaerobiosis. Previously, *dctA* was shown to co-purify with an MS2-tagged AbcR1 transcript in affinity chromatography experiments (Fig. 5C). Supporting these data, computational predictions using IntaRNA indicated that the two previously characterized AbcR1 anti-Shine-Dalgarno regulatory motifs, aSDa (nt 22–32) and aSDb (nt 50–58), may function as seed pairing regions to interact at different sites within the *dctA* CDS, with hybridization energies consistent with productive base-pairing (Fig. 5D). We therefore used pSKi-AbcR1 overexpression to test *dctA* regulation in both wild-type and Δ*rnc* backgrounds (Fig. 5E). Total RNA was extracted from aerobic log-phase cultures of double transconjugants, both before and 16 hours after IPTG induction of wild-type AbcR1 and its variants AbcR1a and AbcR1b (García-Tomsig et al., 2022). These variants carry point mutations in aSDa or aSDb, respectively, which preserve the predicted secondary structure while disrupting base pairing with target mRNAs (Fig. 5E). RT-qPCR revealed that AbcR1 silences *dctA* (expressed constitutively from pR-*dctA*) in the presence of *Sm*RNase III, whereas regulation is lost in the Δ*rnc* mutant, where *dctA* levels increase. Silencing persists with AbcR1b (mutated aSDb) but is abolished with AbcR1a (mutated aSDa), indicating aSDa-mediated regulation. Strikingly, we also observed stronger *dctA* silencing induced by AbcR1b in the wild-type background, likely due to the emergence of a novel interaction site in this mutant variant that forms a duplex with *dctA* more efficiently cleaved by *Sm*RNase III (Fig. S3). Similarly, a new interaction site emerges in the AbcR1a mutant (Fig. S3), which nevertheless does not rescue *Sm*RNase III-dependent silencing. These results confirm that *dctA* mRNA regulation is mediated by the aSDa motif of AbcR1 through a non-canonical mechanism that bypasses the translation initiation region and depends on *Sm*RNase III-mediated decay.

## 4. Discussion

Preserving RNA homeostasis through the activity of ribonucleases and regulatory sRNAs is essential for bacteria to maintain fitness under fluctuating environments. However, understanding of this layer of post-transcriptional gene regulation remains mostly unknown in non-model bacterial species exhibiting complex lifestyles. Among the ribonucleases, RNase III is particularly notable for its ability to recognize and cleave dsRNA regions, thereby contributing not only to mRNA turnover and processing, but also to base-pairing sRNA-mediated silencing. Symbiotic N_2_-fixing rhizobia exemplify versatile bacteria capable of efficiently adapting to shifting environmental conditions, transitioning from a free-living state in soil to an intracellular residence within legume root nodules. Here, we provide genetic evidence of the broad impact of the *S. meliloti* RNase III ortholog, *Sm*RNase III, in maintaining steady-state levels of mRNAs and sRNAs under the microaerobic conditions required for nodule functioning. Our data further show that the proper expression of genes specifying key symbiotic metabolic pathways depends on the coordinated activity of *Sm*RNase III and base-pairing sRNAs, either of the antisense or *trans*-acting classes.

### 4.1. SmRNase III is pivotal in shaping the microaerobic S. meliloti transcriptome

Root nodules that host endosymbiotic bacteroids create a complex environment that drives differential transcription of more than 20% of *S. meliloti* protein-coding genes (Becker et al., 2004). Consequently, strains carrying mutations in genes that pleiotropically affect cell physiology, such as *rnc*, or those involved in core symbiotic functions, typically induce empty or unevenly colonized, non-functional nodules. These defective nodules are unsuitable as biological material for elucidating the molecular mechanisms underlying bacterial-dependent symbiotic phenotypes. A commonly used alternative for investigating symbiotic pathways involves mimicking specific nodule conditions in free-living cultures, where viability of the mutants is uncompromised. In this context, imposing an oxygen downshift on rhizobial cultures, although it does not fully recreate the bacteroid transcriptome, is sufficient to induce expression of a subset of genes involved in bacteroid respiration and symbiotic N_2_ fixation (Becker et al., 2004). Our transcriptomic analyses of the *S. meliloti* wild-type strain reinforced this notion, while revealing substantial changes in both the coding (one-third of *S. meliloti* mRNAs) and non-coding transcriptomes of the *rnc* knockout mutant, consistent with its previously reported pleiotropic effects on free-living and symbiotic phenotypes (Saramago et al., 2018). Under low oxygen conditions, both in culture and within nodule bacteroids, several gene clusters located on the pSymA megaplasmid are notably induced. These include genes involved in rhizobactin biosynthesis, microaerobic denitrification, transcriptional regulation of symbiotic N_2_ fixation, and nitrogenase maturation and function. Data revealed distinct regulation of these pathways by *Sm*RNase III. In the *rnc* knockout mutant, iron-regulated rhizobactin biosynthesis and microaerobic nitrate reduction to N_2_ are downregulated, indicating impaired siderophore production and denitrification. In contrast, transcripts encoding components of the nitrogenase complex are overrepresented, suggesting enhanced expression of the N_2_ fixation machinery. Altered iron acquisition may impair *S. meliloti* competence both in the plant rhizosphere and within nodules, while unbalancing of microaerobic N metabolism compromises symbiotic efficiency. It is also noticeable that exceptionally large sets of oxygen-insensitive genes (1,343 mRNAs and 447 sRNAs) were misregulated in the mutant strain, particularly under microaerobic conditions. This extensive, oxygen-dependent *Sm*RNase III regulon suggests that this rhizobial endoribonuclease has evolved to function efficiently within the nodule environment, playing a pivotal role in buffering transcript levels that support both core cellular processes and genuine symbiotic functions.

### 4.2. SmRNase III substrates inferred from the transcriptomics datasets

Our experimental setup was conceived to assess the global impact of *Sm*RNase III on *S. meliloti* transcriptome, rather than to precisely identify the endogenous substrates of the enzyme. In the mutant strain, upregulated and downregulated transcripts occur in similar numbers compared to the wild type. Although the differential accumulation of many of these RNA species likely reflects indirect regulatory effects, both negative and positive outcomes can be explained by *Sm*RNase III-mediated cleavage at defined sites within its RNA targets, ultimately leading either to RNA degradation or to the production of stable processed transcripts. We therefore scanned sequencing read coverage across all annotated genomic features using a 20-nt sliding window to identify markedly upregulated regions in the *rnc* knockout mutant as most probable *Sm*RNase III cleavage sites. This approach yielded consensus sequence and structural cleavage signatures, identifiable in 28% of mRNAs and 12% of sRNAs influenced by *Sm*RNase III. Notably, a chi-squared test-based statistical analysis revealed that these predicted cleavage sites are enriched in experimentally determined or virtual 5′-UTRs in the mRNAs. Preference for 5′-UTR cleavage has already been reported for *Ec*RNase III in *E. coli*. Examples of conserved mRNAs post-transcriptionally regulated by *Ec*RNase III-mediated processing at the 5′-UTR include *rnc* and *pnp*, which encode *Ec*RNase III itself and polynucleotide phosphorylase (PNPase), respectively, both predicted targets of *Sm*RNase III in *S. meliloti*. In these cases, the regulatory function of the 5′-UTR is to protect the mRNA from RNase E-dependent degradation, such that RNase III-mediated removal of these regions promotes message decay. Interestingly, predictions suggest that 5′-UTRs with more complex post-transcriptional regulatory mechanisms are also substrates of *Sm*RNase III. These include the leader region of thiamine biosynthesis mRNAs (*thi*), which acts as a thiamine-responsive transcriptional attenuator riboswitch in *Rhizobium etli* and *E. coli* (Miranda-Ríos et al., 2001; Rodionov et al., 2002; Guedich et al., 2016). Our pattern-based inference analysis thus provides a valuable resource for interpreting RNase-dependent transcriptional changes assessed by conventional RNA-Seq, while laying a solid foundation for deeper investigation into the mechanisms and regulatory roles of RNase III activity in rhizobia.

### 4.3. SmRNase III assists riboregulation of symbiotic microaerobic pathways

Several reports document the involvement of RNase III in the initial cleavage of RNA duplexes originated through antisense interactions between sRNA regulators and their target transcripts (Svensson and Sharma, 2021; Šetinová et al., 2018). Alterations in the non-coding transcriptome are typically overlooked in conventional RNA-Seq analyses due to misannotations in bacterial genomes. Accurate annotation of the *S. meliloti* genome, including both protein-coding and non-coding loci, therefore enables a more comprehensive and rigorous interpretation of transcriptomic data. Besides mRNA leaders, asRNAs and *trans*-sRNAs were also well represented in our transcriptomic dataset as *Sm*RNase III-dependent RNA species, particularly under microaerobic conditions. The vast majority, if not all, of these sRNAs have unassigned functions. Therefore, our findings position them within a symbiotic context, providing a foundation for reverse genetics-based investigations into their mechanistic principles and regulatory roles.

Massive accumulation of discrete antisense asRNAs in RNase III knockout mutants has been recognized as a hallmark of the pivotal role of this enzyme in mRNA silencing induced by antisense transcription (Lasa et al., 2011). In *S. meliloti*, antisense transcription skews towards pSymA-borne symbiotic genes that are highly expressed in the microaerobic environment of the nodule (Robledo et al., 2015). Of note, we found that the absence of *Sm*RNase III significantly alters the abundance of a substantial number of asRNAs, most prominently upon the oxygen downshift, with more than 90% of these transcripts being upregulated in the mutant. This subset is enriched in asRNAs that target mRNAs encoding both core and accessory components of the nitrogenase enzyme complex, including NifK, NifN, and FixX, which are significantly upregulated in microaerobic cultures of the *rnc* deletion mutant. Expression of several of these asRNAs, including asNifK1 (formerly SmelA031), which is transcribed antisense to the *nifK* mRNA encoding the β subunit of dinitrogenase, had been reliably detected by Northern blot probing of RNA extracted from nodule tissues (Robledo et al., 2015). Our data show that *nifK* and asNifK1 are *Sm*RNase III substrates; however, the antisense interaction enhances *nifK* decay both *in vitro* and *in vivo*. Widespread antisense transcription of N_2_ fixation genes within root nodules may counterbalance the strong microaerobic transcriptional output, helping to maintain the precise stoichiometry of the nitrogenase complex. In this context, our findings underscore a key role for *Sm*RNase III in regulating the biogenesis and assembly of this enzymatic machinery essential for symbiosis.

RNase III has also been associated with post-transcriptional regulation mediated by *trans*-acting sRNAs (Viegas *et al*., 2011). We identified nearly 20% of the *trans*-sRNAs predicted to be expressed by *S. meliloti* as *Sm*RNase III-dependent under microaerobic growth conditions, suggesting that the enzyme may influence their regulatory activity. However, none of these *trans*-sRNAs has undergone functional characterization, and information regarding their potential mRNA targets is currently lacking. To explore potential regulatory connections, we compared the *Sm*RNase III-dependent transcriptome with the metabolic mRNA targetomes of the homologous *trans*-sRNAs AbcR1 and AbcR2, which are the only *trans*-sRNAs in rhizobia for which genome-wide target data are currently available. Interestingly, among the shared mRNA targets of *Sm*RNase III and AbcR1, we identified *dctA*, which encodes a major C4-dicarboxylate transporter. Although AbcR1 expression remains unchanged in the *Sm*RNase III mutant, *dctA* is specifically upregulated under aerobic conditions in the absence of the enzyme. AbcR1/2 are typically expressed in free-living *S. meliloti* and during early stages of host infection to fine-tune metabolic activity, but are absent in the N_2_ fixation zone of mature alfalfa nodules. This expression pattern supports a model in which *Sm*RNase III-dependent silencing of *dctA* occurs endogenously prior to bacteroid differentiation. Our data confirmed that *in vivo* regulation of *dctA* by AbcR1 requires *Sm*RNase III and revealed an uncommon regulatory mechanism involving base pairing deep within the *dctA* coding sequence, mediated by the first (aSDa) of the two known AbcR1 regulatory motifs. This supports the notion that aSDa and aSDb function independently rather than redundantly in target regulation, with each motif controlling distinct sets of target mRNAs (García-Tomsig et al., 2022). Strikingly, both the wild-type AbcR1 transcript and its variants, AbcR1a and AbcR1b, can engage in multiple antisense interactions within *dctA*, some of which even enhance *Sm*RNase III activity. However, only the configuration that preserves wild-type aSDa-*dctA* base-pairing achieves effective silencing. These findings hint at *Sm*RNase III as a novel determinant of AbcR1 target specificity. The spatio-temporal selective silencing of *dctA* by *Sm*RNase III likely fine-tunes *S. meliloti* metabolism during rhizosphere colonization and nodule infection, while enabling dicarboxylate uptake within nodules to meet the high-energy demands of N_2_ fixation.

## 5. Conclusion

Our findings support that *Sm*RNase III is a major regulator of RNA homeostasis in *S. meliloti*, especially under the microaerobic conditions found in root nodules. By controlling extensive coding and non-coding regulons, *Sm*RNase III ensures proper expression of core symbiotic pathways such as iron uptake, denitrification, and nitrogenase assembly. We identified an oxygen-dependent *Sm*RNase III regulon with enriched cleavage signatures in 5′-UTRs, consistent with conserved RNase III targeting mechanisms. *Sm*RNase III also modulates antisense and *trans*-acting sRNAs, influencing both nitrogenase transcript turnover and AbcR1-dependent target regulation. Together, these results position *Sm*RNase III as a pivotal regulator of metabolic and symbiotic functions in the shift of *S. meliloti* from a free-living to a symbiotic lifestyle.

## Acknowledgements

Alicia Barroso (Genomics Unit of Instituto de Parasitología y Biomedicina “López-Neyra”, CSIC, Granada) is acknowledged for RNAseq.

## CRediT authorship contribution statement

**Sabina K. Guedes-García:** Writing – review & editing, Writing – original draft, Visualization, Validation, Methodology, Investigation, Formal analysis, Data curation, Conceptualization. **Natalia I. García-Tomsig:** Writing – review & editing, Writing – original draft, Visualization, Validation, Methodology, Investigation, Formal analysis, Data curation, Conceptualization. **Margarida Saramago:** Writing – review & editing, Validation, Supervision, Resources. **Rute Matos:** Writing – review & editing, Validation, Supervision, Resources. **Cecilia Arraiano:** Writing – review & editing, Validation, Supervision, Resources. **José I. Jiménez-Zurdo:** Writing – review & editing, Writing – original draft, Visualization, Validation, Supervision, Resources, Project administration, Funding acquisition, Formal analysis, Data curation, Conceptualization.

## Data availability

Raw sequence data that support the findings of this study have been deposited in the Sequence Read Archive (SRA) with the primary accession code PRJNA1387402.

## References

Alexa, A., Rahnenführer, J., Lengauer, T., 2006. Improved scoring of functional groups from gene expression data by decorrelating Go graph structure. Bioinformatics 22, 1600–1607. 10.1093/bioinformatics/btl140.

Amarasinghe, A. K., Calin-Jageman, I., Harmouch, A., Sun, W., Nicholson, A. W., 2001. *Escherichia coli* Ribonuclease III: Affinity purification of hexahistidine-tagged enzyme and assays for substrate binding and cleavage. Methods Enzymol. 342, 143–158. 10.1016/s0076-6879(01)42542-0.

Arraiano, C.M., Andrade, J.M., Domingues, S., Guinote, I.B., Malecki, M., Matos, R.G., Moreira, R.N., Pobre, V., Reis, F.P., Saramago, M., Silva, I.J., Viegas, S.C., 2010. The critical role of RNA processing and degradation in the control of gene expression. FEMS Microbiol. Rev. 34, 883–923. 10.1111/j.1574-6976.2010.00242.x.

Au, K.F., Jiang, H., Lin, L., Xing, Y., Wong, W.H., 2010. Detection of splice junctions from paired-end RNA-seq data by SpliceMap. Nucleic. Acids. Res. 38, 4570–4578. 10.1093/nar/gkq211.

Bardwell, J., Régnier, P., Chen, S.M., Nakamura, M., Grunberg-Manago, M., Court, D.L., 1989. Autoregulation of RNase III operon by mRNA processing. EMBO J. 8, 3401–3407. 10.1002/j.1460-2075.1989.tb08504.x.

Batista, M. B., Dixon, R., 2019. Manipulating nitrogen regulation in diazotrophic bacteria for agronomic benefit. Biochem. Soc. Trans. 47, 603–614. 10.1042/bst20180342.

Becker, A., Barnett, M.J., Capela, D., Dondrup, M., Kamp, P.B., Krol, E., Linke, B., Rüberg, S., Runte, K., Schroeder, B.K., Weidner, S., Yurgel, S.N., Batut, J., Long, S.R., Pühler, A., Goesmann, A., 2009. A portal for rhizobial genomes: RhizoGATE integrates a *Sinorhizobium meliloti* genome annotation update with postgenome data. J. Biotechnol. 140, 45–50. 10.1016/j.jbiotec.2008.11.006.

Becker, A., Bergès, H., Krol, E., Bruand, C., Rüberg, S., Capela, D., Lauber, E., Meilhoc, E., Ampe, F., De Bruijn, F.J., Fourment, J., Francez-Charlot, A., Kahn, D., Küster, H., Liebe, C., Pühler, A., Weidner, S., Batut, J., 2004. Global changes in gene expression in *Sinorhizobium meliloti* 1021 under microoxic and symbiotic conditions. Mol. Plant-Microbe Interact. 17, 292–303. 10.1094/mpmi.2004.17.3.292.

Beringer, J.E., 1974. R Factor transfer in *Rhizobium leguminosarum*. Microbiology (N Y) 84, 188–198. 10.1099/00221287-84-1-188.

Del Val, C., Rivas, E., Torres-Quesada, O., Toro, N., Jiménez-Zurdo, J.I., 2007. Identification of differentially expressed small non-coding RNAs in the legume endosymbiont *Sinorhizobium meliloti* by comparative genomics. Mol. Microbiol. 66, 1080–1091. 10.1111/j.1365-2958.2007.05978.x.

Durand, S., Gilet, L., Condon, C., 2012. The essential function of *B. subtilis* RNase III is to silence foreign toxin genes. PLoS Genet. 8, e1003181. 10.1371/journal.pgen.1003181.

Dusha, I., Kondorosi, A., 1993. Genes at different regulatory levels are required for the ammonia control of nodulation in *Rhizobium meliloti*. Mol. Gen. Genet. 240(3), 435–444. 10.1007/bf00280398.

García-Tomsig, N.I., Robledo, M., diCenzo, G.C., Mengoni, A., Millán, V., Peregrina, A., Uceta, A., Jiménez-Zurdo, J.I., 2022. Pervasive RNA regulation of metabolism enhances the root colonization ability of nitrogen-fixing symbiotic α-rhizobia. mBio 13, e03576–21. 10.1128/mbio.03576-21.

Gordon, G.C., Cameron, J.C., Pfleger, B.F., 2017. RNA sequencing identifies new RNase III cleavage sites in *Escherichia coli* and reveals increased regulation of mRNA. mBio 8, e00128. 10.1128/mbio.00128-17.

Guedich, S., Puffer-Enders, B., Baltzinger, M., Hoffmann, G., Da Veiga, C., Jossinet, F., Thore, S., Bec, G., Ennifar, E., Burnouf, D., Dumas, P., 2016. Quantitative and predictive model of kinetic regulation by *E. coli* TPP riboswitches. RNA Biol. 13, 373–390. 10.1080/15476286.2016.1142040.

Hammond, S. M., 2005. Dicing and slicing: the core machinery of the RNA interference pathway. FEBS Letters 579, 5822–5829. 10.1016/j.febslet.2005.08.079.

Huang, L., Deighan, P., Jin, J., Li, Y., Cheung, H., Lee, E., Mo, S.S., Hoover, H., Abubucker, S., Finkel, N., McReynolds, L., Hochschild, A., Lieberman, J., 2020. Tombusvirus p19 captures RNase III-cleaved double-stranded RNAs formed by overlapping sense and antisense transcripts in *Escherichia coli*. mBio 11, e00485. 10.1128/mbio.00485-20.

Ifill, G., Blimkie, T., Lee, A. H., Mackie, G. A., Chen, Q., Stibitz, S., Hancock, R. E. W., Fernandez, R. C., 2021. RNase III and RNase E influence posttranscriptional regulatory networks involved in virulence factor production, metabolism, and regulatory RNA processing in *Bordetella pertussis*. mSphere 6, e0065021. 10.1128/msphere.00650-21.

Jarrige, A., 2001. PNPase autocontrols its expression by degrading a double-stranded structure in the *pnp* mRNA leader. EMBO J. 20, 6845–6855. 10.1093/emboj/20.23.6845.

Jiménez-Zurdo, J.I., Valverde, C., Becker, A., 2013. Insights into the noncoding RNome of nitrogen-fixing endosymbiotic α-proteobacteria. Mol. Plant-Microbe Interact. 26, 160–167. 10.1094/MPMI-07-12-0186-CR.

Jiménez-Zurdo, J.I., Robledo, M., 2015. Unraveling the universe of small RNA regulators in the legume symbiont *Sinorhizobium meliloti*. Symbiosis 67, 43–54. 10.1007/s13199-015-0345-z.

Lasa, I., Toledo-Arana, A., Dobin, A., Villanueva, M., De Los Mozos, I.R., Vergara-Irigaray, M., Segura, V., Fagegaltier, D., Penadés, J.R., Valle, J., Solano, C., Gingeras, T.R., 2011. Genome-wide antisense transcription drives mRNA processing in bacteria. Proc. Natl. Acad. Sci. USA 108, 20172–20177. 10.1073/pnas.1113521108.

Lejars, M., Caillet, J., Solchaga-Flores, E., Guillier, M., Plumbridge, J., Hajnsdorf, E., 2022. Regulatory interplay between RNase III and antisense RNAs in *E. coli*: the case of AsflhD and FlhD, component of the master regulator of motility. mBio 13, 0098122. 10.1128/mbio.00981-22.

Lejars, M., Hajnsdorf, E., 2023. Bacterial RNase III: Targets and physiology. Biochimie 217, 54–65. 10.1016/j.biochi.2023.07.009.

Li, H., Handsaker, R.E., Wysoker, A., Fennell, T., Ruan, J., Homer, N., Marth, G., Abecasis, G.R., Durbin, R., 2009. The sequence alignment/map format and SAMTools. Bioinformatics 25, 2078–2079. 10.1093/bioinformatics/btp352.

Liao, Y., Smyth, G.K., Shi, W., 2019. The R package RSubRead is easier, faster, cheaper and better for alignment and quantification of RNA sequencing reads. Nucleic Acids Res. 47, e47. 10.1093/nar/gkz114.

Love, M.I., Huber, W., Anders, S., 2014. Moderated estimation of fold change and dispersion for RNA-Seq data with DeSeq2. Genome Biol. 15, 550. 10.1186/s13059-014-0550-8.

Lorenz, R., Bernhart, S.H., Siederdissen, C.H.Z., Tafer, H., Flamm, C., Stadler, P.F., Hofacker, I.L., 2011. ViennaRNA Package 2.0. Algorithms Mol. Biol. 6, 26. 10.1186/1748-7188-6-26.

Matsunaga, J., Simons, E.L., Simons, R.W., 1996. RNase III autoregulation: structure and function of rncO, the posttranscriptional “operator”. RNA 2, 1228–1240. https://pubmed.ncbi.nlm.nih.gov/8972772.

Mediati, D.G., Wong, J.M.W., Gao, W., McKellar, S.W., Pang, C.P., Wu, S., Wu, W., Sy, B.M., Monk, I.R., Biazik, J.M., Wilkins, M.R., Howden, B.P., Stinear, T.P., Granneman, S., Tree, J.J., 2022. RNase III-CLASH of multi-drug resistant *Staphylococcus aureus* reveals a regulatory mRNA 3′UTR required for intermediate vancomycin resistance. Nat. Commun. 13, 3558. 10.1038/s41467-022-31177-8.

Mendoza, A., 1995. The enhancement of ammonium assimilation in *Rhizobium etli* prevents nodulation of *Phaseolus vulgaris*. Mol. Plant Microbe-Interact. 8, 584–592. 10.1094/mpmi-8-0584.

Meur, M.R., Portier, C., 1992. *E. coli* polynucleotide phosphorylase expression is autoregulated through an RNase III-dependent mechanism. EMBO J. 11, 2633–2641. 10.1002/j.1460-2075.1992.tb05329.x.

Miranda-Ríos, J., Navarro, M., Soberón, M., 2001. A conserved RNA structure (thi box) is involved in regulation of thiamin biosynthetic gene expression in bacteria. Proc. Natl. Acad. Sci. USA 98, 9736–9741. 10.1073/pnas.161168098.

Oldroyd, G. E., Murray, J. D., Poole, P. S., Downie, J. A., 2011. The rules of engagement in the legume-rhizobial symbiosis. Annu. Rev. Gen. 45, 119–144. 10.1146/annurev-genet-110410-132549.

Patriarca, E. J., Tate, R., Iaccarino, M., 2002. Key role of bacterial NH 4 + metabolism in rhizobium-plant symbiosis. Microbiol. Mol. Biol. Rev. 66, 203–222. 10.1128/mmbr.66.2.203-222.2002.

Persmark, M., Pittman, P., Buyer, J.S., Schwyn, B., Gill, P.R., Neilands, J.B., 1993. Isolation and structure of rhizobactin 1021, a siderophore from the alfalfa symbiont *Rhizobium meliloti* 1021. J. Am. Chem. Soc. 115, 3950–3956. 10.1021/ja00063a014.

Régnier, P., Grunberg-Manago, M., 1990. RNase III cleavages in non-coding leaders of *Escherichia coli* transcripts control mRNA stability and genetic Expression. Biochimie 72, 825–834. 10.1016/0300-9084(90)90192-j.

Robinson, J.A., Thorvaldsdottir, H., Winckler, W., Guttman, M., Lander, E.S., Getz, G., Mesirov, J.P., 2011. Integrative Genomics Viewer. Nat. Biotechnol. 29, 24–26. 10.1038/nbt.1754.

Robledo, M., Frage, B., Wright, P.R., Becker, A., 2015. A stress-induced small RNA modulates alpha-rhizobial cell cycle progression. PLoS Genet. 11, e1005153. 10.1371/journal.pgen.1005153.

Robledo, M., García-Tomsig, N.I., Jiménez-Zurdo, J.I., 2020. Riboregulation in nitrogen-fixing endosymbiotic bacteria. Microorganisms 8, 384. 10.3390/microorganisms8030384.

Rodionov, D.A., Vitreschak, A.G., Mironov, A.A., Gelfand, M.S., 2002. Comparative genomics of thiamin biosynthesis in procaryotes. J. Biol. Chem. 277, 48949–48959. 10.1074/jbc.m208965200.

Saramago, M., Garrido, M., Matos, R.G., Jiménez-Zurdo, J.I., Arraiano, C.M., 2018. *Sinorhizobium meliloti* RNase III: catalytic features and impact on symbiosis. Front. Genet. 9, 350. 10.3389/fgene.2018.00350.

Schlüter, J., Reinkensmeier, J., Daschkey, S., Evguenieva-Hackenberg, E., Janssen, S., Jänicke, S., Becker, J., Giegerich, R., Becker, A., 2010. A genome-wide survey of sRNAs in the symbiotic nitrogen-fixing alpha-proteobacterium *Sinorhizobium meliloti*. BMC Genomics 11, 245. 10.1186/1471-2164-11-245.

Schlüter, J., Reinkensmeier, J., Barnett, M.J., Lang, C., Krol, E., Giegerich, R., Long, S.R., Becker, A., 2013. Global mapping of transcription start sites and promoter motifs in the symbiotic α-proteobacterium *Sinorhizobium meliloti* 1021. BMC Genomics 14, 156. 10.1186/1471-2164-14-156.

Šetinová, D., Šmídová, K., Pohl, P., Musić, I., Bobek, J., 2018. RNase III-binding-mRNAs revealed novel complementary transcripts in *Streptomyces*. Front. Microbiol. 8, 2693. 10.3389/fmicb.2017.02693.

Srivastava, N., Srivastava, R.A.K., 1996. Expression, purification and properties of recombinant *E. coli* ribonuclease III. Biochem. Mol. Biol. Int. 39, 171–180. 10.1080/15216549600201171.

Sun, W., 2005. Catalytic mechanism of *Escherichia coli* ribonuclease III: kinetic and inhibitor evidence for the involvement of two magnesium ions in RNA phosphodiester hydrolysis. Nucleic Acids Res. 33, 807–815. 10.1093/nar/gki197.

Svensson, S.L., Sharma, C.M., 2021. RNase III-mediated processing of a *trans*-acting bacterial sRNA and its *cis*-encoded antagonist. Elife 10, e69064. 10.7554/elife.69064.

Viegas, S.C., Silva, I.J., Saramago, M., Domingues, S., Arraiano, C.M., 2010. Regulation of the small regulatory RNA MicA by ribonuclease III: a target-dependent pathway. Nucleic Acids Res. 39, 2918–2930. 10.1093/nar/gkq1239.

Vogel, J., Argaman, L., Wagner, E.H., Altuvia, S., 2004. The small RNA IstR inhibits synthesis of an SOS-induced toxic peptide. Curr. Biol. 14, 2271–2276. 10.1016/j.cub.2004.12.003.

Wickham, H., 2009. ggplot2: Elegant graphics for data analysis. Springer, New York. ISBN: 978-0-387-98140-6. 10.1007/978-0-387-98141-3.

Yurgel, S. N., Kahn, M. L., 2008. A mutant GlnD nitrogen sensor protein leads to a nitrogen-fixing but ineffective *Sinorhizobium meliloti* symbiosis with alfalfa. Proc. Natl. Acad. Sci. USA 105, 18958–18963. 10.1073/pnas.0808048105.

Zhang, K., Nicholson, A.W., 1997. Regulation of ribonuclease III processing by double-helical sequence antideterminants. Proc. Natl. Acad. Sci. USA 94, 13437–13441. 10.1073/pnas.94.25.13437.

